# Oxytocin reduces top-down control of attention by increasing bottom-up attention allocation to social but not non-social stimuli

**DOI:** 10.1101/519918

**Authors:** Xiaolei Xu, Jialin Li, Zhuo Chen, Keith M. Kendrick, Benjamin Becker

**Author notes:** Correspondence to: Benjamin Becker or Keith M. Kendrick University of Electronic Science and Technology of China The Clinical Hospital of Chengdu Brain Science Institute, MOE Key Laboratory of Neuroinformation No.2006, Xiyuan Ave., West Hi-Tech Zone, Chengdu, Sichuan 611731, China or. equal contributions (Xiaolei Xu, Jialin Li).

## Abstract

The neuropeptide oxytocin (OXT) may facilitate attention to social stimuli by influencing early stage bottom-up processing although findings in relation to different emotional expressions are inconsistent and its influence on top-down cognitive processing mechanisms unclear. In the current double-blind, placebo (PLC) controlled, between subject design study we therefore recruited 71 male subjects to investigate the effects of intranasal OXT (24IU) on both bottom-up attention allocation and top-down attention inhibition using a visual antisaccade paradigm with concurrent eye movement acquisition. Our results show that OXT increased antisaccade errors for social stimuli (all types of emotional faces), but not shapes. This effect of OXT was modulated by trait behavioral inhibition and there was also evidence for reduced state anxiety after OXT treatment. Antisaccades are under volitional control and therefore this indicates that OXT treatment produced reduced top-down inhibition. However, the overall findings are consistent with OXT acting to reduce top-down control of attention as a result of increasing bottom-up early attentional processing of social, but not non-social, stimuli in situations where the two systems are in potential conflict. This effect of OXT is also modulated by individual levels of trait behavioral inhibition possibly as a result of an anxiolytic action.

## 1. Introduction

The neuropeptide oxytocin (OXT) has been repeatedly demonstrated to facilitate early attentional processing of socio-emotional stimuli which may promote facial emotion recognition (Domes et al., 2007; Guastella et al., 2010; Luo et al., 2017). Specifically, the intranasal administration of OXT has been found to increase attention to the eye region (Guastella et al, 2008), to enhance detection of emotions in subliminal presented backward-masked facial stimuli (Schulze et al., 2011) and to augment early attentional orientation towards facial stimuli (Domes et al., 2013; Tollenaar et al., 2013;Xu et al., 2015). Depending on the specific task paradigms employed, OXT has been found to enhance attentional bias towards positive (Domes et al., 2013), positive and neutral (Xu et al., 2015), or positive and negative (Tollenaar et al., 2013) social stimuli. In another study OXT was found to promote switching of attention away from internal interoceptive cues towards external social ones (positive, negative or neutral expression faces) (Yao et al., 2018). Together, these findings suggest a role for OXT in automatic, bottom-up attention processing of salient social-emotional stimuli. However, attention is a strongly limited cognitive resource and efficient allocation of this resource requires a balanced interplay between stimulus-driven bottom-up orientation towards salient stimuli in the environment and top-down goal-directed cognitive control of attention in response to task demands (Buschman and Miller2007; Corbetta et al., 2008). Despite an initial report on the modulatory influence of intranasal OXT on cognitive control during processing of non-social salient stimuli (Striepens et al., 2016), to date differential effects of OXT on bottom-up versus top-down control of attention towards social versus non-social stimuli have not been examined. Against this background the current study combined a randomized double-blind placebo controlled OXT administration protocol with an antisaccade eye-tracking paradigm to determine OXT effects on both bottom-up stimulus-driven attention (prosaccade) and top-down inhibitory control of attention (antisaccade).

Antisaccade paradigms have been widely employed to investigate top-down (volitional) inhibitory control of attention in response to task demands. Briefly, a stimulus is presented in the peripheral region of the visual field and subjects are instructed to either look towards (prosaccade) or away from (antisaccade) the stimulus. The prosaccade eye gaze represents a stimulus-driven reflexive behavioral response towards a potentially salient stimulus in the environment. In contrast, a successful antisaccade requires the initial inhibition of the stimulus-driven automatic prosaccade as well as the subsequent volitional saccade away from the distractors, reflecting a top-down inhibitory attentional control mechanism (Munoz and Everling, 2004).

In view of frequently reported social-specific effects of OXT (Shamay-Tsoory and Abu-Akel, 2016) the present study employed a social-emotional antisaccade paradigm including neutral, happy, sad, fearful and angry facial expressions as social and oval shapes as non-social stimuli to determine whether OXT generally modulates attention allocation or specifically modulates the processing of social stimuli. Moreover, given previous reports on valence-specific effects of OXT on processing of social stimuli (Xu et al., 2015), stimuli of different facial emotions were included to allowed to further explore emotion-specific effects of OXT on social attention allocation. Given that overarching hypotheses on the modulatory influence of OXT on social cognition suggest that its augmentation of social salience represents a core mechanism of action across different functional domains (Shamay-Tsoory and Abu-Akel, 2016) we hypothesized that OXT would specifically increase attentional bias for social stimuli as reflected by facilitated prosaccades in the context of impaired inhibition of volitional attentional control (impaired antisaccades). Based on our previous findings suggesting that OXT specifically enhances attention allocation towards neutral and positive facial stimuli (Domes et al., 2013; Xu et al., 2015) we further hypothesized that OXT would particularly affect processing of neutral and positive (happy) faces rather than that of negative ones such as sad, fearful, and angry faces.

## 2. Materials and methods

### 2.1 Participants

71 healthy male students aged 18-30 years (mean ± sem = 21.85 ± 0.32 years) from the University of Electronic Science and Technology of China (UESTC) were recruited for the present randomized, placebo-controlled, double blind between-subject pharmaco-eye tracking study. Exclusion criteria were any previous or current neurological or psychiatric disorders, as well as current (30 days before the experiment) or regular use of any psychotropic substances, including nicotine. All participants were instructed to abstain from alcohol and caffeine during the 24 hours before the pharmacological eye-tracking experiment. Participants were randomly assigned to receive either 24 International Units (IU) of intranasal OXT (n = 34, mean ± sem age = 21.88 ± 0.44 years) or placebo (PLC, n = 37, mean ± sem age = 21.81 ± 0.46 years).

Study protocols had full approval by the local ethics committee at the UESTC and experimental in accordance with the latest revision of the Declaration of Helsinki. All participants provided informed consent before the experiment and received monetary compensation for participation. Study protocols and primary outcomes were pre-registered at clinical trials.gov (https://clinicaltrials.gov/ct2/show/NCT03486925, Trial ID: NCT03486925).

### 2.2 Experimental protocols and procedures

To control for confounding effects of between-group differences in variables that have previously been demonstrated to modulate the effects of intranasal OXT (Kendrick et al., 2017) potential confounders were assessed before treatment administration by means of validated scales. Based on previous findings suggesting that individual variations in these variables modulate the effects of OXT, the following variables were assessed: childhood maltreatment (Childhood Trauma Questionnaire, CTQ) (Bernstein and Fink, 1998), social anxiety (Liebowitz Social Anxiety Scale, LSAS; Social Interaction Anxiety Scale, SIAS) (Mattick and Clarke, 1998; Heimberg et al., 1999), state anxiety (State-Trait Anxiety Inventory, STAI) (Barnes et al., 2002), depressive symptom load (Beck Depression Inventory, BDI-II) (Beck et al., 1996), autism traits (the Adult Autism Spectrum Quotient, ASQ) (Baron-Cohen et al., 2001) and mood (Positive and Negative Affect Schedule, PANAS) (Watson and Clark, 1988). In line with the focus of the study on cognitive control towards emotional stimuli, additional scales included the Action Control Scale (ACS) (Kuhl, 1994), Behavioral Inhibition System and Behavioral Activation System Scale (BIS/BAS) (Charles and White, 1994) and Emotion Regulation Questionnaire (ERQ) (Wang et al., 2015). Next, participants self-administrated either 24 IU of OXT (Oxytocin-spray, Sichuan Meike Pharmaceutical Co., Ltd; 3 puffs of 4IU per nostril with 30s between each puff) or PLC (PLC – identical sprays with the same ingredients other than OXT). Administration adhered to standardized intranasal OXT protocols (Guastella et al., 2013), and in line with the pharmacodynamics of intranasal OXT in humans (Spengler et al., 2017) treatment was administered 45 minutes before the eye-tracking paradigm. Mood (PANAS) and state anxiety (STAI-S) were additionally assessed after the experiment to control for unspecific effects of treatment on these domains.

Individual differences in the behavioral inhibition system (BIS) have been associated with both, attentional processing and top-down control of orientation towards social-emotional stimuli, including antisaccade performance (Dennis and Chen, 2009; Mogg et al., 2012). To determine whether OXT affects this association, correlations between individual variations in behavioral inhibition and eye gaze indices that showed OXT effects were further examined within the treatment groups.

### 2.3 Antisaccade paradigm

We employed a modified antisaccade paradigm (Chen et al., 2014) that included 5 social-emotional conditions (facial expressions from 4 male and 4 female actors: neutral, happy, fearful, sad, angry expressions) and a non-social control condition (oval shape, the shape was slightly varied to create 8 different shape stimuli). A total of 576 trials over 14 blocks were presented including 2 non-social control blocks (one anti- and one pro-saccade block) and 12 emotional blocks (6 anti- and 6 pro-saccade blocks). Each emotional block contained 40 trials in randomized order including 8 trials per emotional condition resulting in 48 trials in total per anti- and pro-saccade condition respectively. To avoid carry-over of emotion-specific effects of OXT the paradigm started with the shape blocks, which also contained 48 trials, followed by the emotional blocks. The order of anti- and pro-saccade blocks was randomized. Each block started with a 2000ms visual instruction (“Towards”, “Away”) indicating whether the subsequent block required a pro- or an anti-saccade response followed by a jittered fixation cross (1000-3500ms; mean duration = 1500ms). Following the fixation period a stimulus was presented at 8° visual angle relative to the centered fixation cross to the left or right visual field for 1000ms. The size of stimuli was 400×500 pixels. Participants were asked to direct their gaze as fast as possible towards the stimulus during the prosaccade (“Towards”) blocks and away from it in the opposite direction during the antisaccade blocks (“Away”) (**Fig 1**).

**Figure 1.**
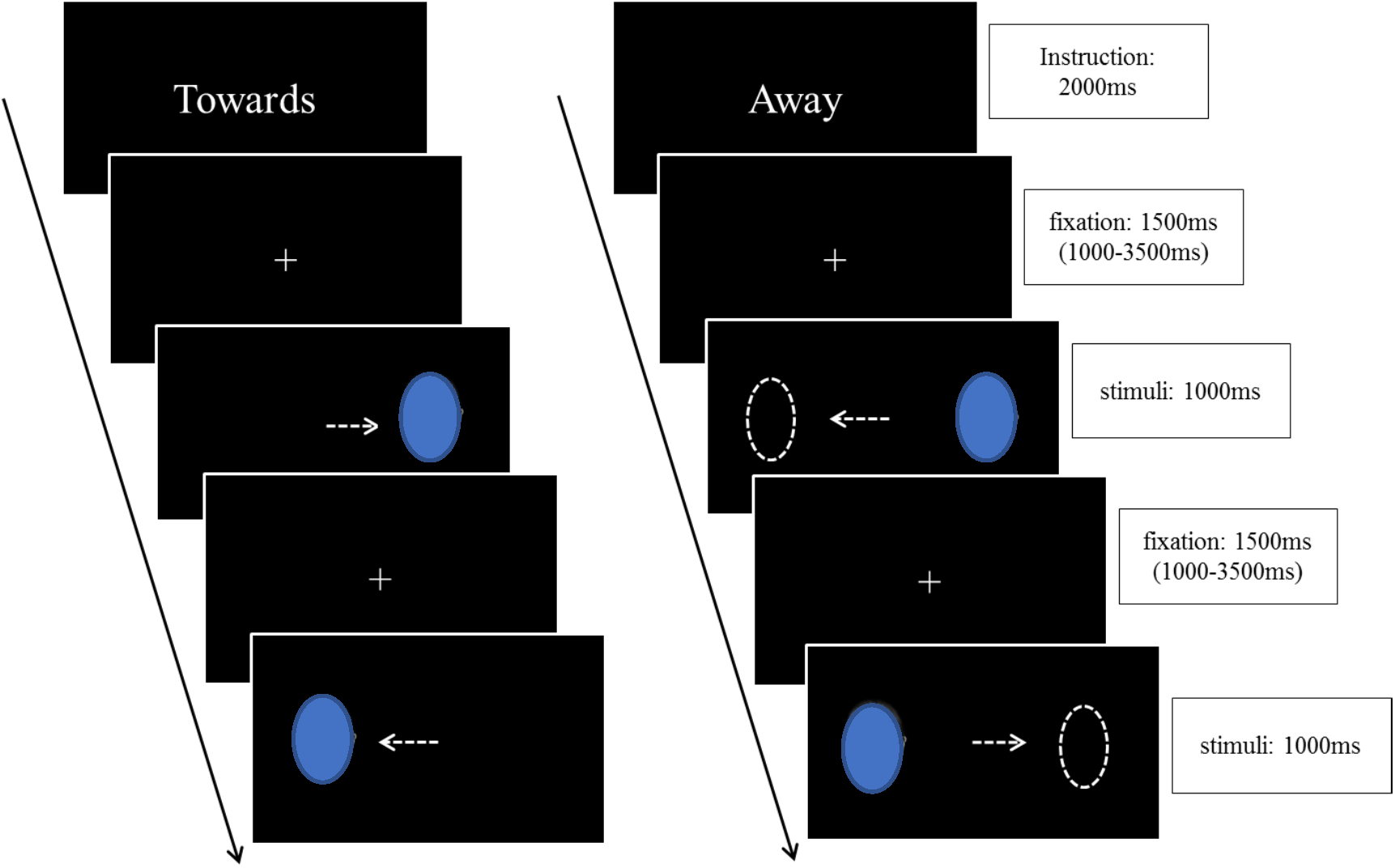
Each block started with an instruction indicating whether a prosaccade (“Towards”) or an antisaccade (“Away”) response was required, followed by a fixation cross centered on the screen. A stimulus was presented at the left or right peripheral position after the fixation disappeared. For “Towards” blocks, subjects were asked to look at the stimulus (prosaccade) and for “Away” blocks they were instructed to look away from the stimulus to the opposite position (antisaccade).

Subjects completed the experiment in a dimly illuminated room. Stimuli were presented on a 17inch monitor at a resolution of 1024×768 pixels. A chin rest was used to standardize the distance and position from the screen (57cm away and centrally positioned relative to the monitor). The eye gaze data was acquired using an EyeLink 1000 Plus system (SR Research, Ottawa, Canada) in monocular mode at a sampling rate of 2000Hz. Before each block a 9-point calibration was conducted, the experimental blocks were divided by brief rest periods to facilitate attentive processing throughout the experiment. For the subsequent analyses saccades were excluded based on the criteria of amplitude < 1.5°, velocity < 30°/sec, and latencies shorter than 80ms which were classified as anticipatory saccades (for a similar strategy see Wieser et al., 2009). The number of excluded anticipatory saccades did not differ between the treatment groups (p = 0.23). The raw eye gaze data was initially exported and processed using the EyeLink DataViewer 3.1 (SR Research, Mississauga, Ontario, Canada) and effects of treatment were subsequently analyzed using SPSS 18.0 software (SPSS Inc., Chicago, Illinois, USA).

Mean error rates for pro- and anti-saccade blocks and latencies for correct saccades served as primary outcomes to determine effects of treatment. Mixed ANOVA models and independent sample t-tests were employed to determine differences between the treatment groups. Post-hoc tests incorporated Bonferroni correction for multiple comparisons. Associations between individual variations in pre-treatment behavioral inhibition (BIS score) and eye gaze behavior were examined using Pearson correlation and differences in the correlations between the two treatment groups were assessed by Fisher’ Z test with appropriate Bonferroni correction.

## 3. Results

### 3.1 Potential confounders

Examining the pre-treatment data on potential confounders using independent t-tests did not reveal significant differences between the two treatment groups (details see Table 1). Likewise, no significant post-treatment differences between the treatment groups were observed for mood. Together, these findings argue against potential confounding effects of pre-treatment differences or unspecific treatment effects. While there were no significant post-treatment differences between the PLC and OXT groups for state-anxiety (*p* = 0.07) there was a significant reduction in state anxiety in the OXT group when comparing pre vs post-treatment scores (*p* = 0.027), but not in the PLC group (*p* = 0.882, **Fig 2**). This suggests that OXT treatment produced anxiolytic effects within the group treated with OXT.

**Table 1.**
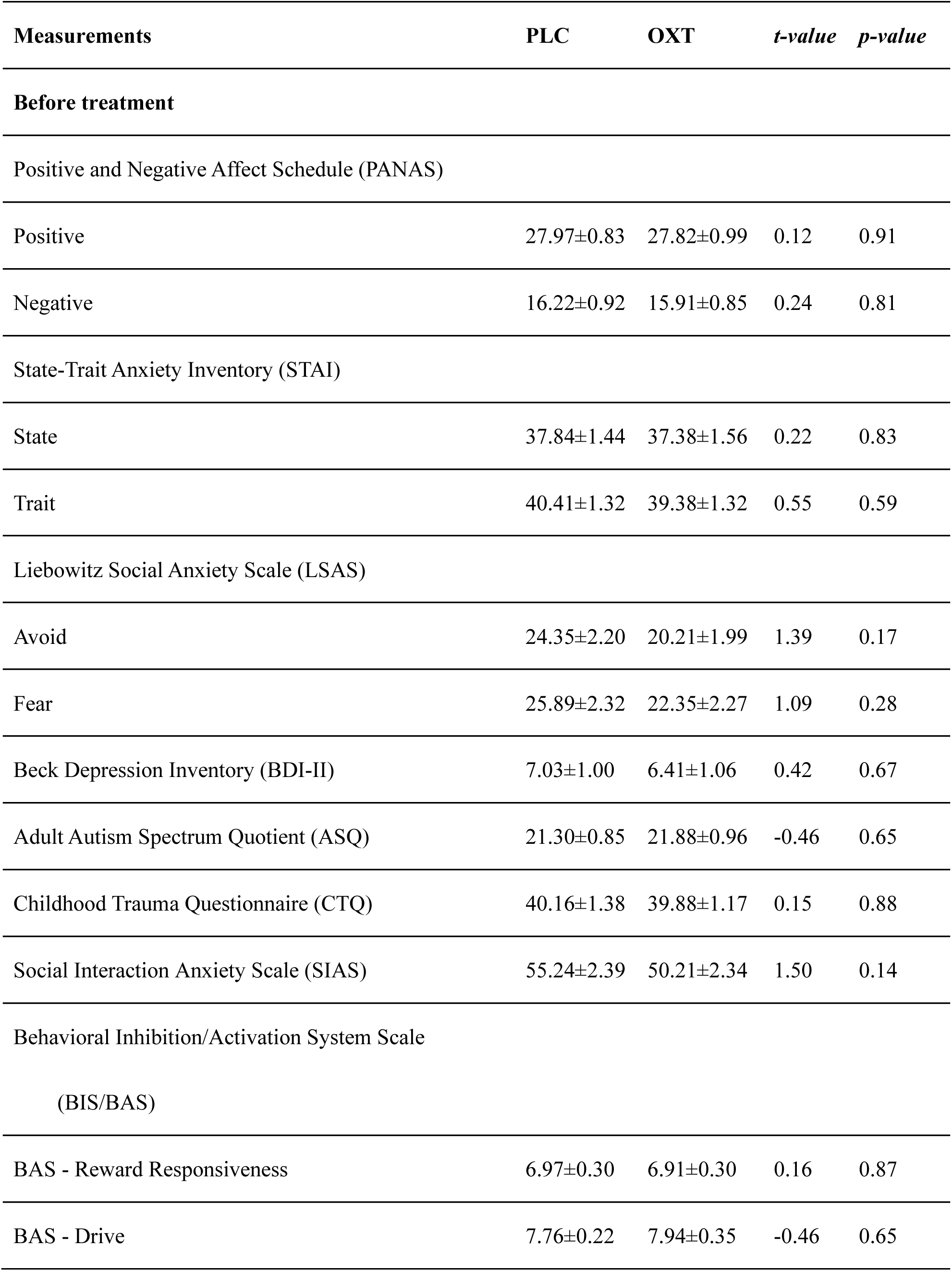
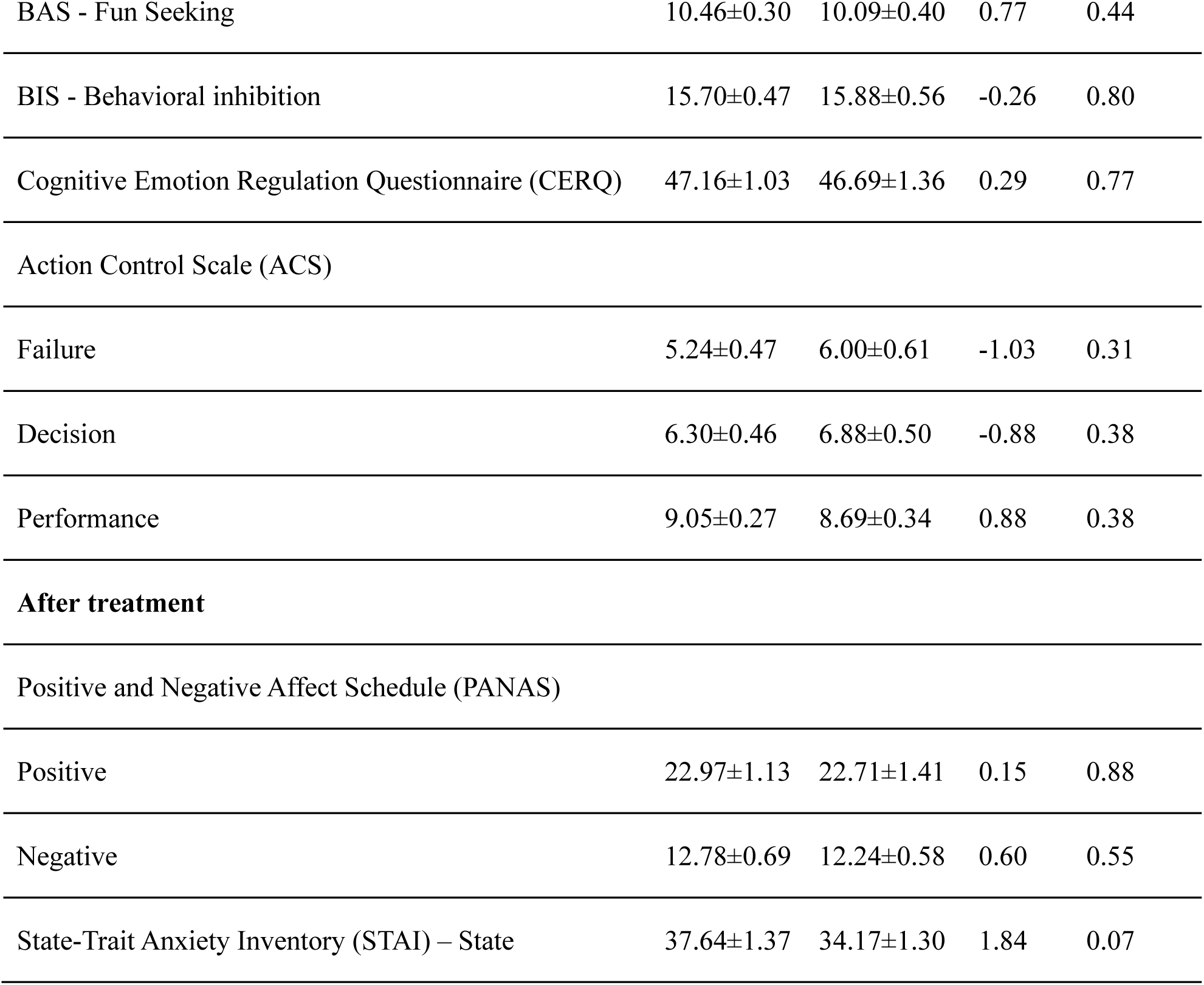
Questionnaire scores in PLC and OXT groups before and after treatment

**Figure 2.**
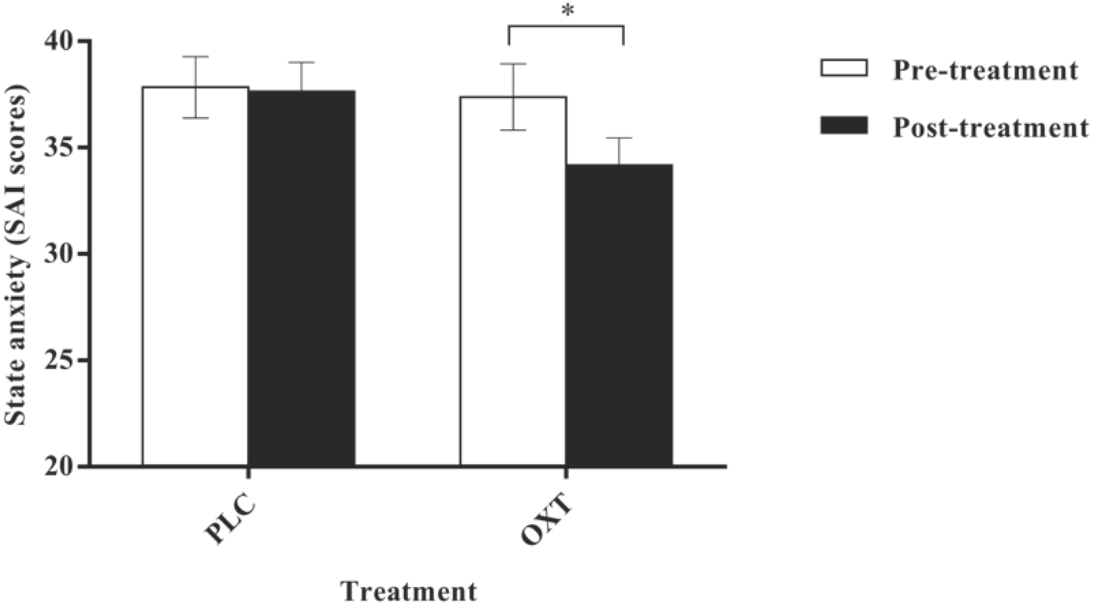
State anxiety (SAI) scores assessed by State-Trait Anxiety Inventory (STAI) before and after treatment. OXT significantly decreased state anxiety after the experiment within the oxytocin-treatment group. Abbreviations: PLC, placebo, OXT, oxytocin

### 3.2 Eye-tracking data

Eye movement data from 4 subjects were excluded based on the quality assessment criteria (see methods, also Wieser et al., 2009) leading to a final sample of PLC = 33 and OXT = 34 for the eye gaze data analyses.

#### 3.2.1 Saccade latency

A 2 (treatment: OXT/PLC) × 2 (stimuli: non-social shape/social-face) × 2 (task: anti-/pro-saccade) mixed ANOVA was conducted to examine saccade latencies. Results revealed significant main effects of task (*F*_1,65_ = 819.57, *p*<0.001, η^2^_p_ = 0.93) and stimuli (*F*_1,65_ =77.14, *p*<0.001, η^2^ _p_= 0.54), as well as a significant stimuli × treatment (*F*_1,65_ = 4.66, *p*=0.035, η^2^_p_ = 0.07) and stimuli × task (*F*_1,65_ = 64.84, *p*<0.001, η^2^_p_ = 0.50) interaction effects. Post-hoc Bonferroni-corrected paired comparisons indicated that the antisaccade latency was significantly longer than the prosaccade one (antisaccade: mean±sem = 277.71±3.33ms, prosaccade: mean±sem = 192.74±2.04ms, *p*<0.001, Cohen’s *d* = 3.23) and that saccades for social-face stimuli were generally faster than those for shape stimuli (non-social, shape, mean±sem = 242.98±2.74ms; social, face stimuli, mean±sem = 227.47±2.21ms, *p*<0.001, Cohen’s *d* = 0.31). Examining the stimuli × treatment interaction effect revealed that saccades for the social-face stimuli were faster in both treatment groups (PLC: non-social, shape, mean±sem = 245.83±3.91ms, social, face stimuli, mean±sem = 226.51±3.15ms, *p*<0.001, Cohen’s *d* = 0.82; OXT: non-social, shape, mean±sem = 240.12±3.85ms, social, face stimuli, mean±sem = 228.43±3.11ms, *p*<0.001, Cohen’s *d* = 0.49; **Fig 3**). Specifically, longer saccade latencies for the shapes were observed during prosaccade but not antisaccade blocks (antisaccade: non-social, shape = 279.28±3.74ms, social, face stimuli = 276.13±3.47ms; *p* = 0.258, Cohen’s *d* = 0.10; prosaccade: non-social, shape = 206.67±2.60ms, social, face stimuli = 178.81±1.78ms, *p*<0,001, Cohen’s *d* = 1.55). Together the results indicate that task instruction (pro-versus anti-saccade) and stimuli (social vs non-social) successfully modulated the behavioral response during the paradigm.

**Figure 3.**
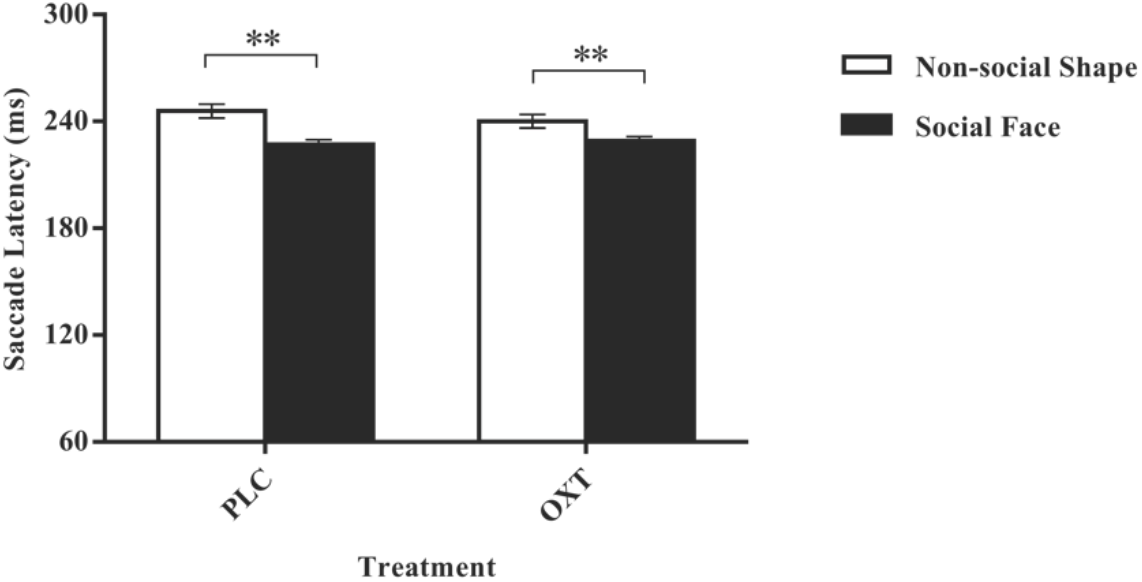
Saccade latencies for shape and face stimuli in the placebo and oxytocin treated group respectively. Abbreviations: PLC, placebo, OXT, oxytocin

### 3.2.2 Saccade error rate

A 2 (treatment: OXT/PLC) × 2 (stimuli: non-social shape/social-face) × 2 (task: anti-/pro-saccade) mixed ANOVA was carried out including saccade error rates as dependent variable. Results revealed significant main effects of stimuli (*F*_1,65_ = 21.87, *p*<0.001, η^2^_p_ = 0.25), task (*F*_1,65_ = 112.16, *p*<0.001, η^2^_p_ = 0.63), and treatment (*F*_1,65_ = 8.88, *p* = 0.004, η^2^_p_ = 0.12) as well as significant task × treatment (*F*_1,65_ = 4.67, *p* = 0.034, η^2^ _p_ = 0.067) and task × stimuli (*F*_1,65_ = 8.15, *p* = 0.006, η^2^ _p_ = 0.11) interaction effects. Post-hoc Bonferroni-corrected comparisons demonstrated that the error rate for shapes was significantly lower compared to social-face stimuli (non-social, shape: mean±sem = 15.53%±1.19, social, face stimuli: mean±sem = 20.40%±1.20, *p*<0.001, Cohen’s *d* = 0.39) and the error rate for antisaccades was significantly higher compared to prosaccades (antisaccade: mean±sem = 22.92%±1.25%, prosaccade: mean±sem = 13.01%±1.09, *p*<0.001, Cohen’s *d* = 0.84). Overall, OXT significantly increased the error rate compared to PLC (PLC: mean±sem = 14.75%±1.54, OXT: mean±sem = 21.18%±1.51, *p* = 0.004, Cohen’s *d* = 0.59). In addition, there was a marginal significant three-way treatment × stimuli × task interaction effect (*F*_1,65_ = 3.03, *p* = 0.086, η^2^_p_ = 0.045) and exploratory analyses revealed that OXT specifically increased error rates for antisaccades of social face stimuli (antisaccade: social, face stimuli: PLC = 20.78%±2.09, OXT = 32.35%±2.06, *t* = 3.94, *p*<0.001, Cohen’s *d* = 0.97, **Fig 4a**).

**Figure 4.**
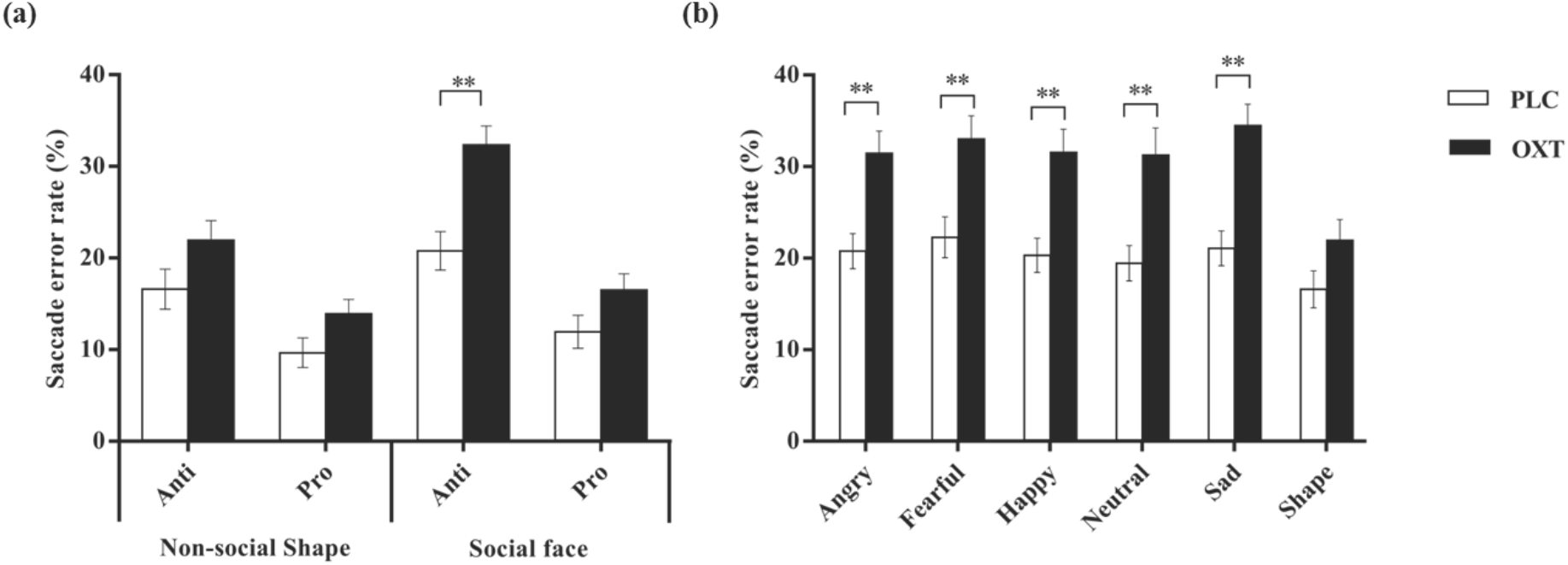
(a) Oxytocin increases the error rate in the social face but not the shape condition in antisaccade blocks; (b) Oxytocin generally increases the error rate for each face emotion but not for shapes during antisaccade blocks. \*\**p*<0.005 Abbreviations: PLC, placebo, OXT, oxytocin

To further explore potential emotion-specific effects of OXT on antisaccade performance an additional 2 (treatment: OXT/PLC) × 6 (stimuli-emotions: shape/happy/angry/neutral/sad/fear) mixed ANOVA with antisaccade error rates as dependent variable was performed. Results revealed a main effect of stimuli (*F*_5,325_ = 10.52, *p*<0.001, η^2^_p_ = 0.14, error rate of shape is significantly lower than all emotional conditions, all *p*s<0.02) and treatment (*F*_1,65_ = 15.30, *p*<0.001, η^2^_p_ = 0.19) suggesting that OXT significantly increased antisaccade error rates compared to PLC (PLC = 20.09%±1.92, OXT = 30.62%±1.89, Cohen’s *d* = 0.80, *p*<0.001). A marginal significant drug × stimuli interaction effect (*F*_1,325_ = 2.01, *p* = 0.077, η^2^_p_ = 0.03) and post-hoc exploratory analysis further revealed that OXT significantly increased the error rates for the all social stimuli but not the shape stimuli (OXT vs. PLC, for all face emotion conditions *p*s<0.002, shape *p*=0.09, effect size for the social-emotion specific post-hoc comparisons: angry: Cohen’s *d* = 0.83; fearful: Cohen’s *d* = 0.78; happy: Cohen’s *d* = 0.87; neutral: Cohen’s *d* = 0.81; sad: Cohen’s *d* = 1.09; shape: Cohen’s *d* = 0.43, **Fig 4b)**.

### 3.3 Effects of oxytocin on the associations between eye gaze and trait behavioral inhibition

To assess modulatory effects of OXT on the association between individual differences in pre-treatment behavioral inhibition and cognitive control of attention, Pearson correlations between trait inhibition (BIS score) and antisaccade error rates were examined separately in the treatment groups. Correlation analyses revealed that the antisaccade error rate for social stimuli was significantly positively associated with pre-treatment BIS scores following OXT but not PLC (PLC: r = -0.189, *p* = 0.292, OXT: r = 0.397, *p* = 0.022, correlation difference, Fisher’s Z = -2.368, *p* = 0.018, Cohen’s *q* = 0.27, **Fig 5**). This suggests that OXT particularly blunted the cognitive inhibition of stimulus-driven attention towards social stimuli in subjects with higher levels of trait behavioral inhibition.

**Figure 5.**
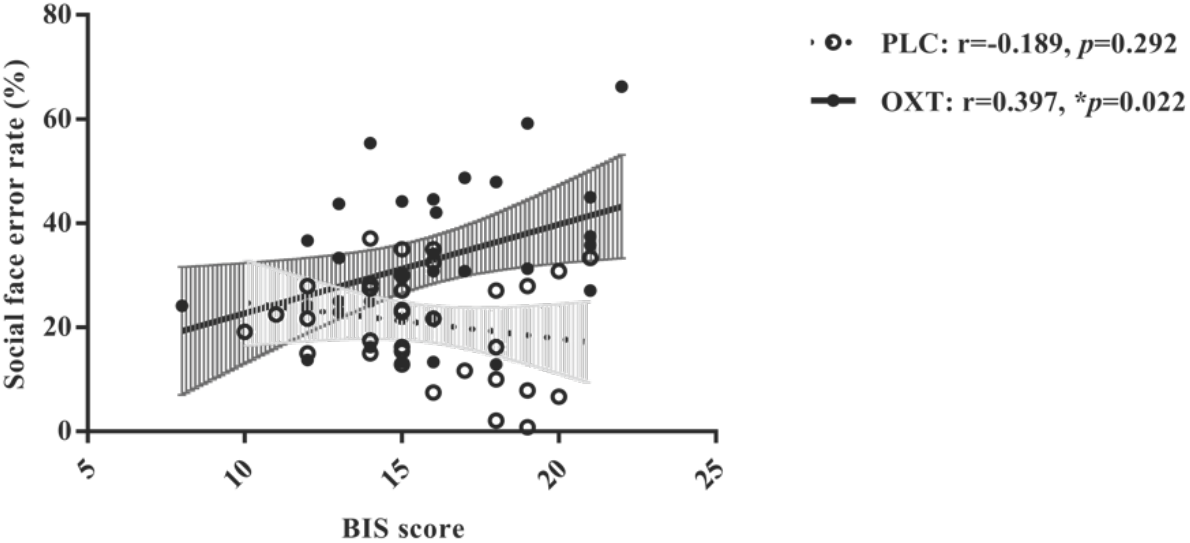
Trait behavioral inhibition as assessed by the BIS scale was positively associated with the error rates for social stimuli antisaccades following OXT but not in PLC treatment. \**p*<0.05 Abbreviations: PLC, placebo, OXT, oxytocin

## 4. Discussion

Overall, results from the present pharmacological eye-gaze study employing a social-emotional pro- and anti-saccade paradigm revealed that participants respond to emotional faces faster than to shapes in the prosaccade blocks and are more prone to errors in the antisaccade blocks, indicating that social stimuli capture greater attention and are more difficult to inhibit responses towards. Furthermore, OXT significantly increased the error rate for facial stimuli during the antisaccade but not the prosaccade condition, suggesting that it decreases the ability to switch attention away from social stimuli, but not non-social stimuli compared with PLC treatment. The effect was significantly positively associated with trait behavioral inhibition as assessed by BIS scores in the OXT, but not the PLC group indicating that OXT is particularly decreasing the ability of subjects with higher general levels of behavioral inhibition to switch their attention away from social but not non-social stimuli.

Our finding that OXT decreased the ability to switch attention away from emotional faces but not shapes is in line with previous studies reporting that it specifically increased attention towards social (neutral and positive expression faces) but not non-social stimuli in an attentional-blink task (Xu et al., 2015) and enhanced attention towards distracting external social stimuli (all face expressions) in an interoceptive awareness task (Yao et al., 2018). Studies have increasingly demonstrated that OXT plays an important role in attention orientation and attention regulation to social cues (for review: Shamay-Tsoory and Abu-Akel, 2016). Rapid and accurate recognition of others’ emotions from their facial expressions is critical for human social interactions (Adolphs, 2002) and OXT has been found to enhance this ability e.g. by increasing attention to the eye region of human faces (Domes et al., 2007; Guastella et al., 2008). The antisaccade gaze requires volitional inhibition of automatic attention allocation towards sudden-onset visual targets and antisaccade performance is strongly modulated by the salience of the stimulus (Myers et al., 2011). The social salience hypothesis of OXT has proposed that OXT is of particular importance for regulating attention towards salient social cues (Shamay-Tsoory and Abu-Akel, 2016). This enhanced social salience effect of OXT might therefore have contributed to an increased difficulty in inhibiting saccades away from facial, but not shape, stimuli, resulting in higher error rates for antisaccadic eye-gaze behavior in the context of social stimuli.

Previous studies investigating the effects of OXT on attentional bias towards specific emotions revealed inconsistent findings. Some studies reported that OXT specifically enhanced attention allocation towards neutral or positive facial stimuli (Domes et al., 2013; Xu et al., 2015) while other studies found that it reduced attentional bias towards negative emotion ones (Kim et al., 2014), increased attention orientation to faces expressing either positive and negative emotions (Tollenaar et al., 2013) or all faces irrespective of emotion (Yao et al., 2018). Although we hypothesized on the basis of these previous findings indicating that OXT can influence early attentional processing of salient social stimuli via acting on bottom-up processing, we failed in the current study to find supportive evidence in terms of either reduced latencies or errors rates when subjects made reflexive prosaccades towards face stimuli. One early study on OXT modulation of detection of social stimuli also failed to find evidence for effects on early perceptual stage processing using a visual search task (Guastella et al., 2009). While this might be considered as evidence for the absence of effects of OXT on bottom-up processing it is notable that in the current task prosaccade errors were very few (13%) and latencies very fast as one would expect for a reflexive response. As such, this might represent a ceiling effect leading to a limited sensitivity of the prosaccade condition to capture OXT treatment effects on bottom-up social attention allocation.

The robust effect of OXT (>50% increase across face emotions, effect size Cohen’s *d*= 0.78-1.09) on increasing antisaccade errors could either be interpreted as evidence for it selectively weakening top-down control processing, without influencing bottom-up control, or alternatively as it increasing bottom-up reflexive mechanisms resulting in impaired top-down control. While OXT only influenced antisaccade and not prosaccade errors, the net effect of this is that subjects under OXT made more prosaccades towards social stimuli so this can be considered as evidence of increased bottom-up processing. On the other hand, top-down attention inhibition is not biased by specific emotions and the effects of OXT on antisaccade errors occurred across all the face emotions. It is clear from previous studies that across different paradigms that OXT often influences bottom-up processing of specific face emotions (Domes et al., 2013; Kim et al., 2013; Tolenaar et al., 2013; Xu et al., 2015) although one study has shown OXT effects irrespective of the specific face emotion (Yao et al., 2018). Interestingly, the latter study showing effects of OXT on bottom up processing across all face emotions was the only one where the paradigm involved potential conflict between top-down and bottom up processing and as such is similar to the antisaccade paradigm. Thus, it is possible that OXT does primarily influence bottom-up attentional processing of salient social stimuli such as faces but whether it alters attention towards specific face expressions or all of them may depend on the extent to which cognitive top-down control and bottom-up reflexive processing mechanisms are in conflict.

The behavioral inhibition system inhibits movements towards targets which may lead to negative outcomes and increase negative feelings such as anxiety and nervousness (Carver and White, 1994). In the current study we found that subjects with higher behavioral inhibition as measured by the BIS showed pronounced effects of OXT with respect to increased antisaccade errors. Individuals with higher BIS scores tend also to exhibit higher social anxiety (Kimbrel et al., 2012) which is associated with increasing antisaccade error rates in response to all facial expressions (Wieser et al., 2009). While OXT has generally been associated with anxiolytic effects (see Kendrick et al., 2017) several studies have also reported that it can have anxiogenic ones (Striepens et al., 2012; Grillon et al., 2013). In the present study there was however evidence for a significant decrease rather than increase in state anxiety following OXT, but not PLC, treatment. Thus, it is possible that OXT reduced social anxiety in subjects with higher BIS scores resulting in them being less motivated to avoid looking at social stimuli which in turn may have increased antisaccade error-rates for social stimuli.

In conclusion, the current study has demonstrated that OXT may primarily function to increase bottom-up processing of attention to salient social stimuli and that where there is a context of conflicting top-down cognitive processing the influence of the latter over bottom-up processing is weakened, leading to more errors in the cognitive task component. The influence of OXT in this context is modulated by trait behavioral inhibition, possibly as a result of its anxiolytic effects.

